# In Vitro Inhibition of SARS-CoV-2 Infection by Bromhexine hydrochloride

**DOI:** 10.1101/2022.12.23.521817

**Authors:** Ronaldo Martins, Iasmin Ferreira, Daniel M. M. Jorge, Leticia Almeida, Juliano P. Souza, Marjorie Pontelli, Italo A. Castro, Thais M. Lima, Rosa M. M. Viana, Dario Zamboni, Priscyla D. Marcato, Eurico Arruda

**Author notes:** **Corresponding author:** Ronaldo Martins; Department of Cell Biology, University of São Paulo School of Medicine, Av. Bandeirantes 3900, Ribeirão Preto, SP, Brazil, 14049-900.

## Abstract

The world is enduring the SARS CoV-2 pandemic, and although extensive research has been conducted on the issue, only a few antivirals have been approved to treat patients with COVID-19. Bromhexine hydrochloride was previously identified as a potent inhibitor of TMPRSS2, an essential protease for ACE-2 virus receptor interactions. In the present study, we investigated whether bromhexine treatment could reduce SARS CoV-2 replication in vitro. To evaluate bromhexine’s effectiveness against SARS COV-2 infection, viral load was measured using Caco-2 cell lines expressing TMPRSS2. Our molecular docking results indicate that bromhexine displays an affinity with the active site of TMPRSS2. The drug significantly inhibited SARS CoV-2, both parental and P1 variant strains, infection in the Caco-2 cell line, reducing about 40% of SARS-CoV-2 entrance and about 90% of viral progeny in the supernatant 48h post-infection. Furthermore, bromhexine did not exhibit any direct virucidal activity on SARS CoV-2. In conclusion, bromhexine hydrochloride efficiently disrupts SARS CoV-2 infection in vitro and has the potential to become an effective antiviral agent in COVID-19 treatment.

## 1. Introduction

Since the beginning of 2020, the world has been overwhelmed by a pandemic of a novel Coronavirus named severe acute respiratory syndrome coronavirus-2 (SARS-CoV-2), which causes coronavirus disease-19 (COVID-19). While numerous studies have been carried out to discover SARS CoV-2 inhibitory drugs, only a few antiviral compounds have been repurposed, and no specific effective anti-SARS-CoV-2 agent has reached clinical use. The best studied SARS CoV-2 cell entry mechanism depends on the binding of the viral spike (S) protein to the angiotensin-converting enzyme 2 (ACE-2) on the cell surface, which is preceded by the priming cleavage of S by host cell membrane-associated type II transmembrane serine protease (TMPRSS2) (Hoffmann et al., 2020). Thus, several groups have proposed repurposing compounds that interfere with TMPRSS, like bromhexine hydrochloride (BHH), to treat COVID-19 patients (Maggio & Corsini, 2020; Fu et al., 2020; Habtemariam et al., 2020). BHH selectively inhibits TMPRSS2 activity (Lucas et al., 2014) and, consequently, could suppress the cleavage of SARS-CoV-2 S protein, limiting the reproduction of the virus. The present study aimed to analyze the in vitro antiviral activity of BHH for SARS CoV-2.

We performed an in silico docking prediction to evaluate the interaction between TMPRSS2 and BHH, followed by in vitro biological assays using the original and the gamma (P1) variants of SARS-CoV-2. Our pre-clinical experiments revealed that BHH significantly inhibits SARS-CoV-2 infection and replication in vitro in a susceptible human cell line that expresses TMPRSS2 in a dose-dependent manner.

## 2. Material and methods

### 2.1. Viruses and compounds

The SARS-CoV-2 Brazil/SPBR-02/2020, and SARS CoV-2 gamma (P1) strain MAN87209 variants were used for *in vitro* experiments. In BSL3 conditions, Vero cells (ATCC CCL-81) were used for virus propagation by inoculation in serum-free Dulbecco’s minimum essential medium (DMEM) supplemented with Penicillin (10,000 U/mL), Streptomycin (10,000 μg/mL) and trypsin-TPCK (1 μg/μL) for 48 hours at 37°C in 5% CO_2_. When cytopathic effect (CPE) was observed, monolayers were harvested with a cell scraper, snap-frozen in liquid nitrogen, thawed and clarified by centrifugation (1000 x g), and the supernatants were aliquoted and stored at −80°C. Later, virus titration was performed in Vero CCL81 cells by serial decimal dilutions to determine the 50% tissue culture infectious dose (TCID_50_) of the virus stocks (Reed & Muench, 1938; Harcourt et al., 2020). Bromhexine hydrochloride (BRH) (Shanghai Shengxin Medicine Chemical Co., Ltd.) was kindly donated by Ourofino; Camostat mesylate were purchased from Sigma Aldrich; Dulbecco’s modified Eagle medium (DMEM) and fetal bovine serum (FBS) were purchased from Gibco (USA); Neutral red dye and DMSO, from Merck.

### 2.2. One-Step Real-Time PCR

SARS-CoV-2 genome quantification was conducted with primer-probe sets for N_2_ and the RNAse-P housekeeping gene, following USA-CDC protocols (Table 1) (Lu et al., 2020). We did One-step real-time RT-PCR (Step-One Plus real-time PCR thermocycler, Applied Biosystems, Foster City, CA, USA), using total nucleic acids extracted by Trizol® (Invitrogen, CA, USA), according to the manufacturer’s instructions. Briefly, 100 ng of RNA was used for genome amplification, adding specific primers (20 μM), probe (5 μM), and TaqPath 1-Step qRT-PCR Master Mix (Applied Biosystems, Foster City, CA, USA), with the following cycling parameters: 25°C for 2 min, 50°C for 15 min, and 95°C for 2 min, followed by 45 cycles of 95°C for 3 s and 55°C for 30s. To assess the SARS-CoV-2 viral load, a standard curve was constructed using plasmids containing the insert. A 944-bp amplicon was inserted into a TA cloning vector (PTZ57R/T CloneJetTM Cloning Kit Thermo Fisher®), starting from residue 14 of the N gene, which includes all 3 sets of primers/probe proposed by the CDC (N1, N2, and N3). For virus genome quantification, serial 10-fold dilutions of the plasmid were prepared, and the SARS-CoV-2 RNA copies were plotted using the GraphPad® Prism 8.4.2 software.

**Table 1.**
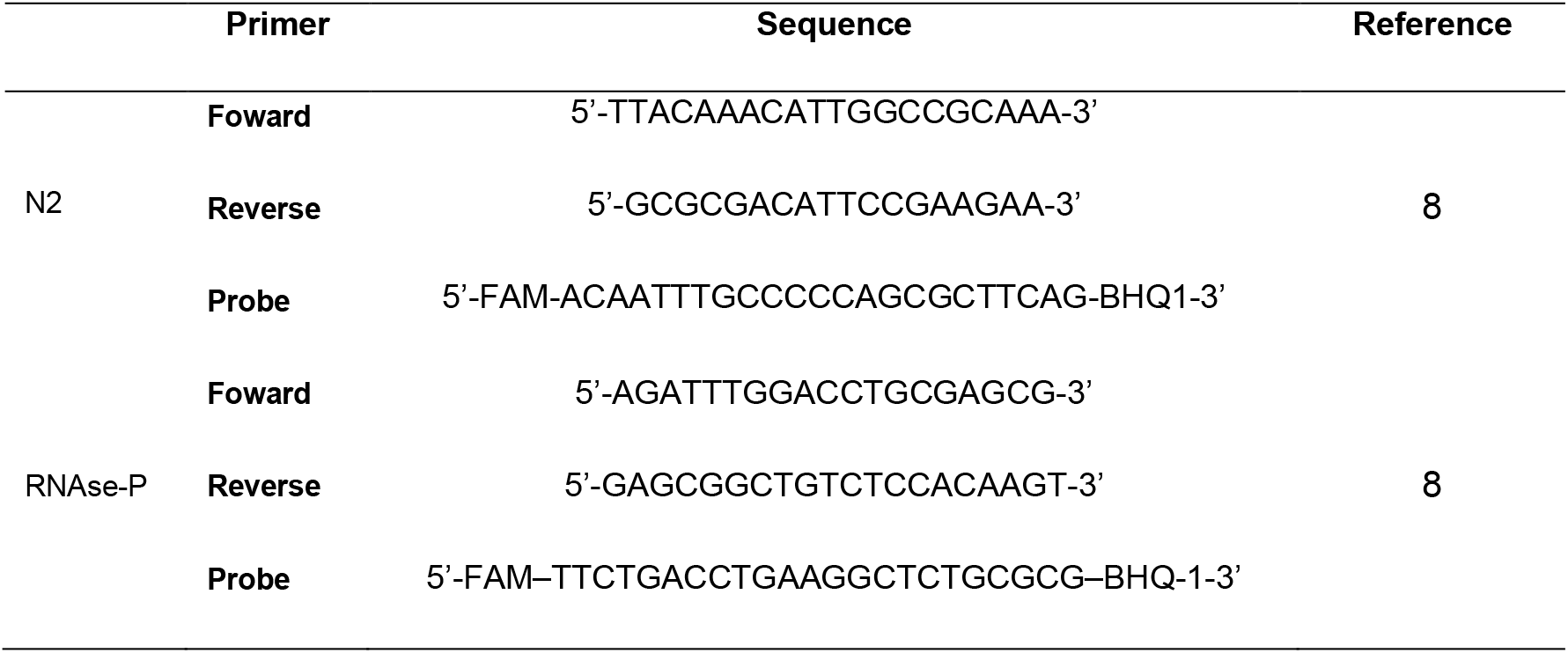
Primer/probe sequences for the detection of SARS-CoV-2 genome and the RNAse-P housekeeping gene.

### 2.3. Homology Modeling and molecular docking analysis of human TMPRSS2

The transmembrane serine protease TMPRSS2 (accession number: O15393) was obtained from the Uniprot database (Nucleic Acids Res, 2019). The TMPRSS2 extracellular domain was used to build a homology model using the SWISS-MODEL web server (Waterhouse et al, 2018), and the serine protease hepsin (protein data bank id: 5ce1.1.A) was selected as a template to predict the three-dimensional structure. The sequence similarity between TMPRSS2 and the serine protease hepsin was 0.38, with coverage of 0.70. The final model was refined using ModRefiner (Xu & Zhang, 2011) and evaluated with PROSA (Wiederstein et al., 2007) and PROCHECK (Laskowski et al., 1993). The refined homology model of the protein TMPRSS2 was used for the molecular docking analysis. Bromhexine hydrochloride (CID 5702220) was obtained from the PubChem Database (https://pubchem.ncbi.nlm.nih.gov/) in sdf format and then converted to mol2 format. The molecular docking of the protein TMPRSS2 and the bromhexine hydrochloride molecule was performed using the Achilles blind docking server (https://bio-hpc.ucam.edu/achilles/). The active site of the TMPRSS2 protein which contains the amino acids His^296^, Asp^345^, and Ser^441^ (Wilson et al., 2005), was used in the blind docking assay to identify the possible sites for bromhexine interaction. The BHH-TMPRSS2 interactions were visualized with AutoDock (Pettersen et al., 2004) and UCSF Chimera (Morris et al., 2009).

### 2.4. Cytotoxicity assay

The cytotoxicity of BHH in Caco-2 cells (Human, colon - ATCC HTB-37) was determined by the neutral red (NR) and Deoxyribonucleic Acid (DNA) Content assays (Correa, 2005; Borefreund & Puerner, 1984; Miranda et al., 2019). The cells were seeded at 5×10^4^ cells/well in 96-well plates. After 24 h, they were treated with different BHH concentrations (0.75 – 200 μM) for 48 h. The negative control was treated only with medium, and the positive control, with 1% DMSO. For the cell viability NR assay, after specific treatment cells were washed once with PBS and incubated for 3 h with serum-free medium containing NR (50 μmol/L). Then, the cells were washed twice with PBS-Ca^+2^, followed by addition of 0.1 mL of 1% acetic acid:50% ethanol. The absorbance of each well was read at 540 nm in a microplate reader SpectraMax i3 (Molecular Devices). After the NR assay, the plate was air-dried, and the DNA content assay was done. For cell lysis, 0.1 mL of 0.5 M NaOH was added and then the cell culture plates were kept at 37°C for 1h. The absorbance was read at 260 nm in a spectrophotometer (Implen, Munich, Germany). The IC_50_ values were calculated by nonlinear regression using the GraphPad® Prism 8.4.2 software.

### 2.5. Inhibition of infection by VSV-eGFP-SARS-CoV-2-SΔ21 by BHH

To assess the effect of BHH to inhibit the viral entry in the intestinal human cell line Caco-2 (ATCC, HTB-37) we used a pseudotyped fluorescent infectious vesicular stomatitis virus able to encode the SARS-CoV-2 S protein in place of the native VSV envelope glycoprotein (VSV-eGFP-SARS-CoV-2-SΔ21) (Case et al., 2020). Briefly, 5×10^4^ Caco-2 cells/well were seeded in 96-well black tissue culture plates and infected with VSV-eGFP-SARS-CoV-2-SΔ21 at MOI=1 in the presence of BHH at 250, 500, 750 and 1000 nM, or vehicle (0.004% DMSO). At 2 hours post-inoculation, the cells were washed with PBS and replenished with fresh medium containing BHH or vehicle at the same concentrations. After 48 h post-infection, images were acquired using ImageXpress Micro XLS System® (Molecular Devices, LLC, EUA), using 6 randomly chosen fields at 40x magnification. For image acquisition the filters used were λex:377 and λem:447 for nuclear DAPI and λex:490 and λem:525 for VSV-eGFP-SARS-CoV-2-SΔ21. The mean cell numbers and fluorescence per cell were obtained using the algorithm Transfluor from MetaXpress Software (version 5.3.0.5, Molecular Devices, LLC). For nuclei counting, the algorithm was set to detect structures between 5-15 μm and intensity of 1200 gray levels above background. For VSV-eGFP-SARS-CoV-2-SΔ21 fluorescence quantification, the settings were chosen to detect structures between 1-2 μm and intensity of 500 gray levels above background. Data were processed using GraphPad® Prism 8.4.2 software and expressed as average of the six fields acquired.

### 2.5. Virucidal effect of BHH on SARS COV-2

To evaluate the direct virucidal activity of BHH on SARS-CoV-2, 5×10^6^ TCID_50_ of SARS-CoV-2 was incubated with BHH diluted in serum-free DMEM at a final concentration of 1500 nM. Two hours later, virus titers were determined by end-point titration by TCID_50_ in 96-well plates.

### 2.6. Antiviral activity assay

The anti-SARS CoV-2 activity of BHH was investigated in vitro using the cell lineages Caco-2 (human colon, ATCC HTB-37), HUH7 (human liver, JCRB0403) and Vero (monkey kidney, ATCC CCL-81). 24-well plates treated with the compounds were infected in triplicate with SARS-CoV-2 (MOI=1), and incubated for adsorption for 2 hours at 37°C in 5% CO_2_. The inoculum was then removed, cultures were washed with PBS, and then treated with medium containing 750 nM BHH or vehicle (0.004% DMSO). The positive control consisted of 100 mM camostat mesylate (Sigma Aldrich), a known TMPRSS2 inhibitor. After 48 h, 250 μL of supernatant was collected for RNA extraction and viral load quantification by real-time RT-PCR using a standard curve as previously reported (Lu et al., 2020). Next an in vitro dose-response experiment was done with Caco-2 cells pretreated with BHH ranging from 250 to 1500 nM for 2 hours, followed by inoculation with SARS-CoV-2 (MOI=1) for 2 h. The monolayers were then washed with PBS and replenished with fresh medium containing BHH at the same concentrations used in the pretreatment. Supernatants were collected for viral genome quantification 48 hours post-infection.

### 2.7. Statistical significance

Statistical significance was determined by ordinary one-way ANOVA, followed by Bonferroni’s post hoc test. Differences were considered statistically significant when p < 0.05. Statistical analyses and graph plots were carried out using the GraphPad® Prism 8.4.2 software.

## 3. Results

### 3.1. Binding of BHH to TMPRSS2

Molecular docking showed that bromhexine binds to TMPRSS2. The initial search was conducted based on the 492 amino acids of TMPRSS2, and the hits obtained in the homology modeling mapped to a portion of the extracellular domain. A three-dimensional structure of this domain, from position 146 to 489, was predicted using the serine protease hepsin (5ce1.1.A). The PRoSA web server revealed that the final model had a z-score of −8.8 (PROCHECK tool), indicating that 96.78% of the residues were present in the allowed regions. The MolProbity tool yielded a value of 2.36, with an overall qmean −2.11, which was higher than the initial model, a z-score of −8.76, and 92.20% of residues were present in the allowed regions, MolProbity of 1.89 and an overall quality qmean of −1.47. The final predicted model was used in the docking analysis since the binding site for bromhexine was unknown. A blind docking server was also used to generate a three-dimensional structure representing the interaction between TMPRSS2 and BHH, with 15 possible clusters of binding poses. The selected bromhexine pose in the active site of TMPRSS2 exhibited a −5.7 kcal/mole binding energy, with hydrogen bonds with Ser460 (Fig. 1).

**Figure 1.**
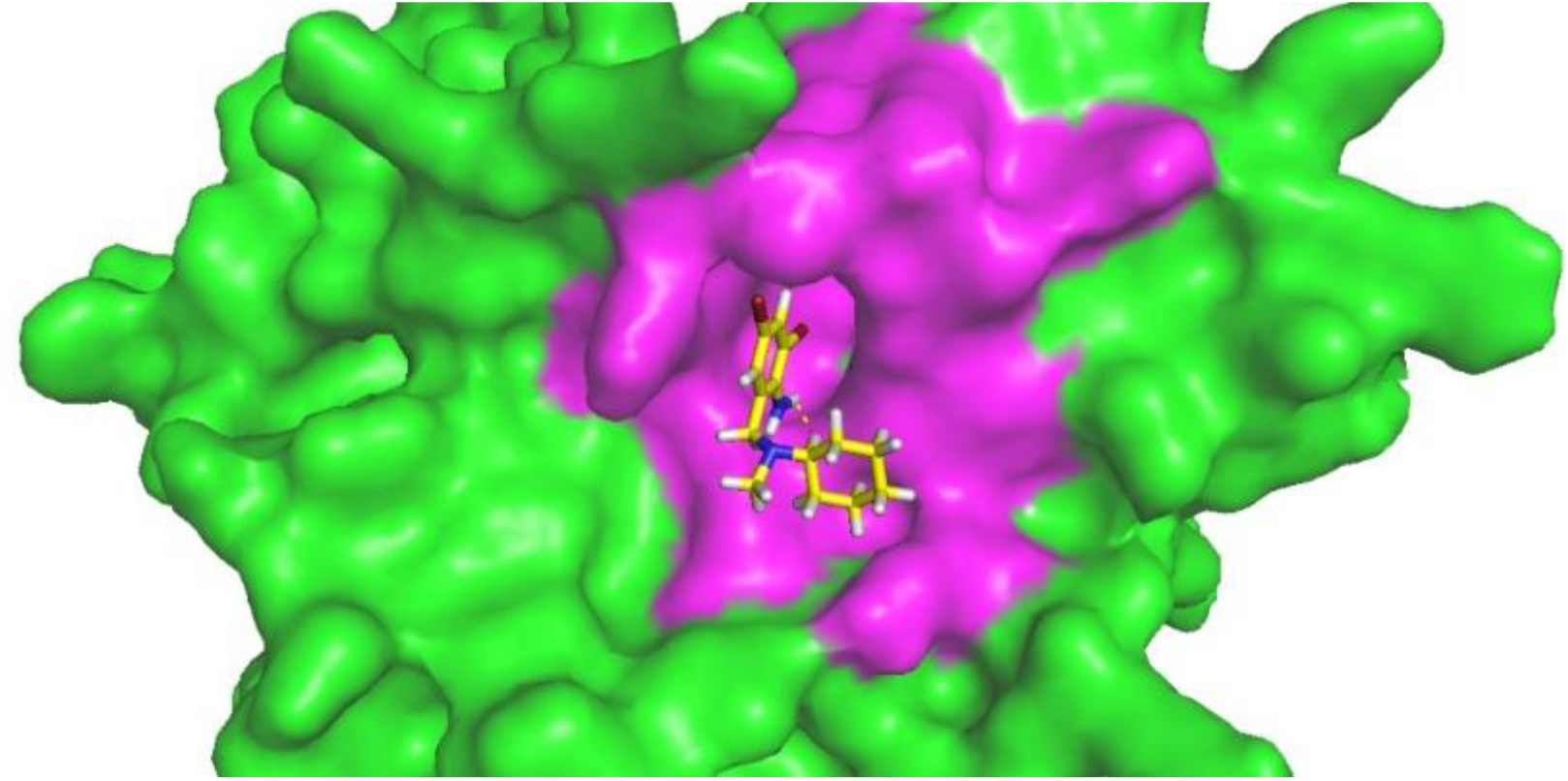
Docking model using the Achilles blind docking server between the 3D structures of the TMPRSS2 protein and BHH. Green and pink colors indicate the TMPRSS2 protein and its active site residue, respectively. The chemical elements represent BHH.

### 3.2. In vitro cytotoxicity

A dose-dependent cytotoxicity effect of bromhexine on Caco-2 cells was observed after 48h of exposure, as shown in Figure 2. Cell viability was maintained at 100% in bromhexine concentrations ≤25 μM, and the IC_50_ was 76.52 μM. However, cell viability above 70% was observed with bromhexine concentration up to 50 μM (Fig. 2).

**Figure 2.**
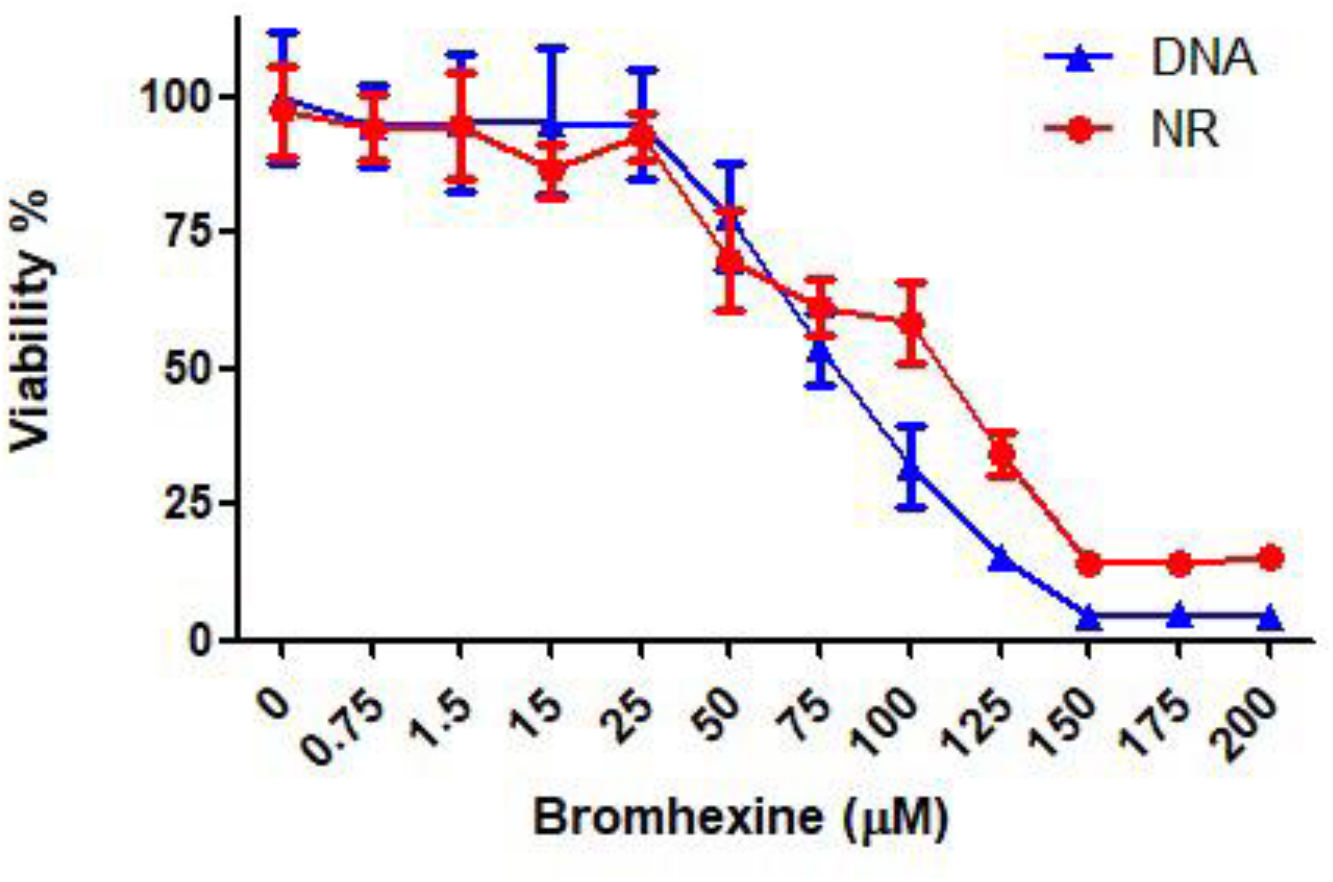
Viability of Caco-2 cells after treatment with bromhexine for 48 h. The Neutral Red (NR) uptake and nucleic acid content (DNA) were determined in triplicate. Each point represents the mean±S.D.

### 3.3. Inhibition of VSV-eGFP-SARS-CoV-2-SΔ21 with BHH

BHH was able to impair the infection of recombinant VSV expressing S protein of SARS-CoV-2 using Caco-2 cells. Images acquired 48 h post-infection using a fluorescence microscope indicated that BHH treatment lead to reduction of GFP expression during VSV-eGFP-SARS-CoV-2-SΔ21 replication in Caco-2 cells of approximately of 40% at 500nM concentration of BHH (Fig. 3).

**Fig 3.**
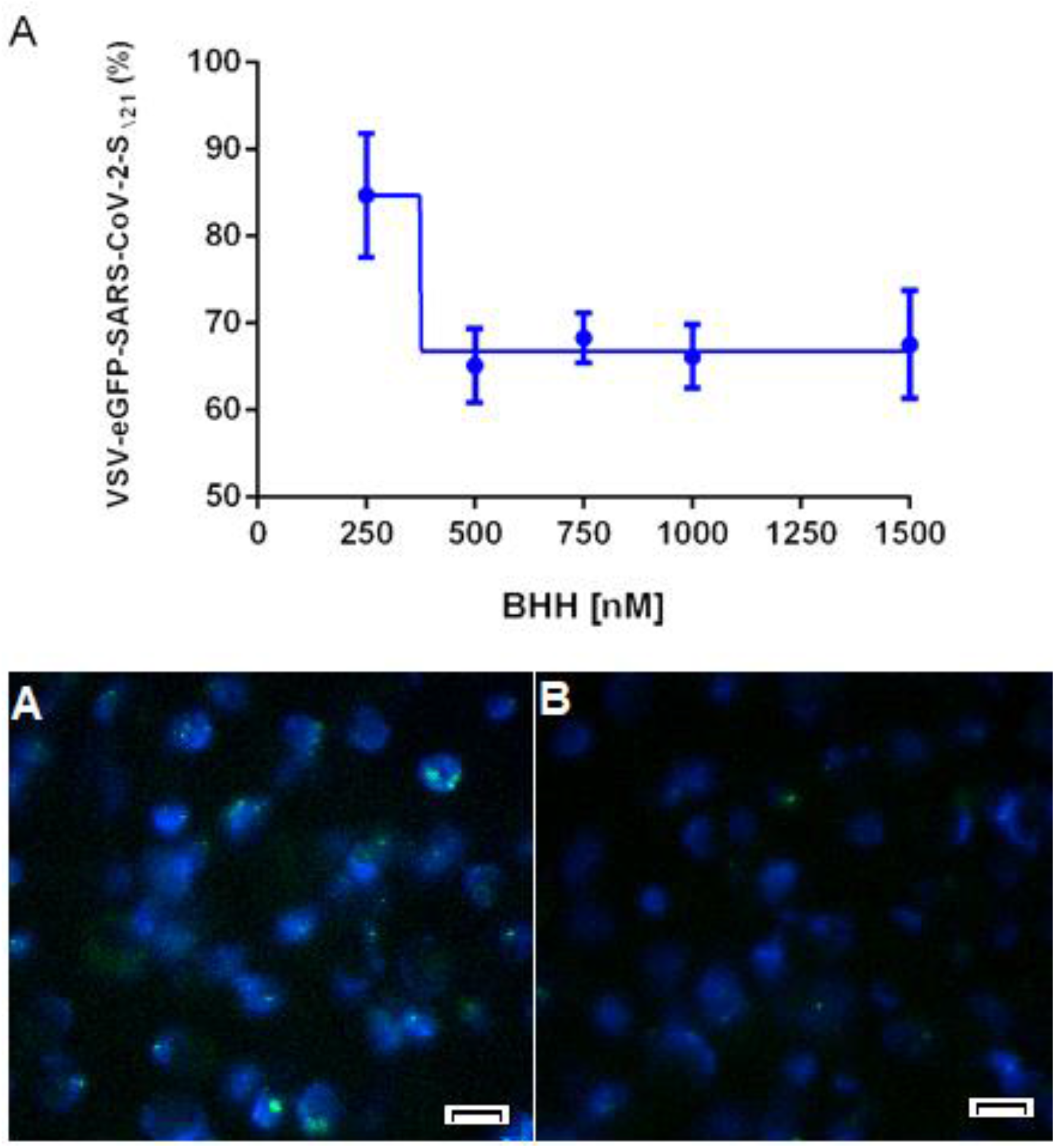
Inhibition of VSV-eGFP-SARS-CoV-2-SΔ21 in Caco-2 cells by BHH. (A) Percentual BHH inhibition of VSV-eGFP-SARS-CoV-2-SΔ21 replication in Caco-2 cells 48 h post-infection. (B-C) Caco-2 cells infected with VSV-eGFP-SARS-CoV-2-SΔ21 (green) and treated with 0.004% DMSO (B) or 1.5 μM of BHH (C). Nuclei stained in blue. Scale bar 85 μm.

### 3.4. In vitro antiviral effect

When we tested the antiviral activity of bromhexine using a non-cytotoxic concentration (0.75 μM), we verified that the drug strongly inhibited SARS-CoV-2 infection in Caco-2 cells (p < 0.001) (Fig. 4A). Remarkably, bromhexine was more effective in reducing the SARS-CoV-2 replication than the positive control camostat mesylate (p < 0,0001), with 1.5-fold-higher inhibition efficiency than camostat mesylate. Additionally, a dose-response effect of bromhexine on SARS-CoV-2 replication *in vitro* was observed in the Caco-2 cells infected with both parental and P1 variant strains. The estimated inhibitory concentration (IC_50_) of bromhexine was around 1μM (Fig. 4B). Moreover, antiviral effects of bromhexine over SARS-CoV-2 infection have not been reproduced when we use HUH7 and Vero CCL-81 cells lines. Infectious virus titration experiments determined by TCID_50_ revealed that the drug did not exhibit direct virucidal activity at 1500 nM after 2 hours of incubation at room temperature (Fig. 5).

**Fig 4.**
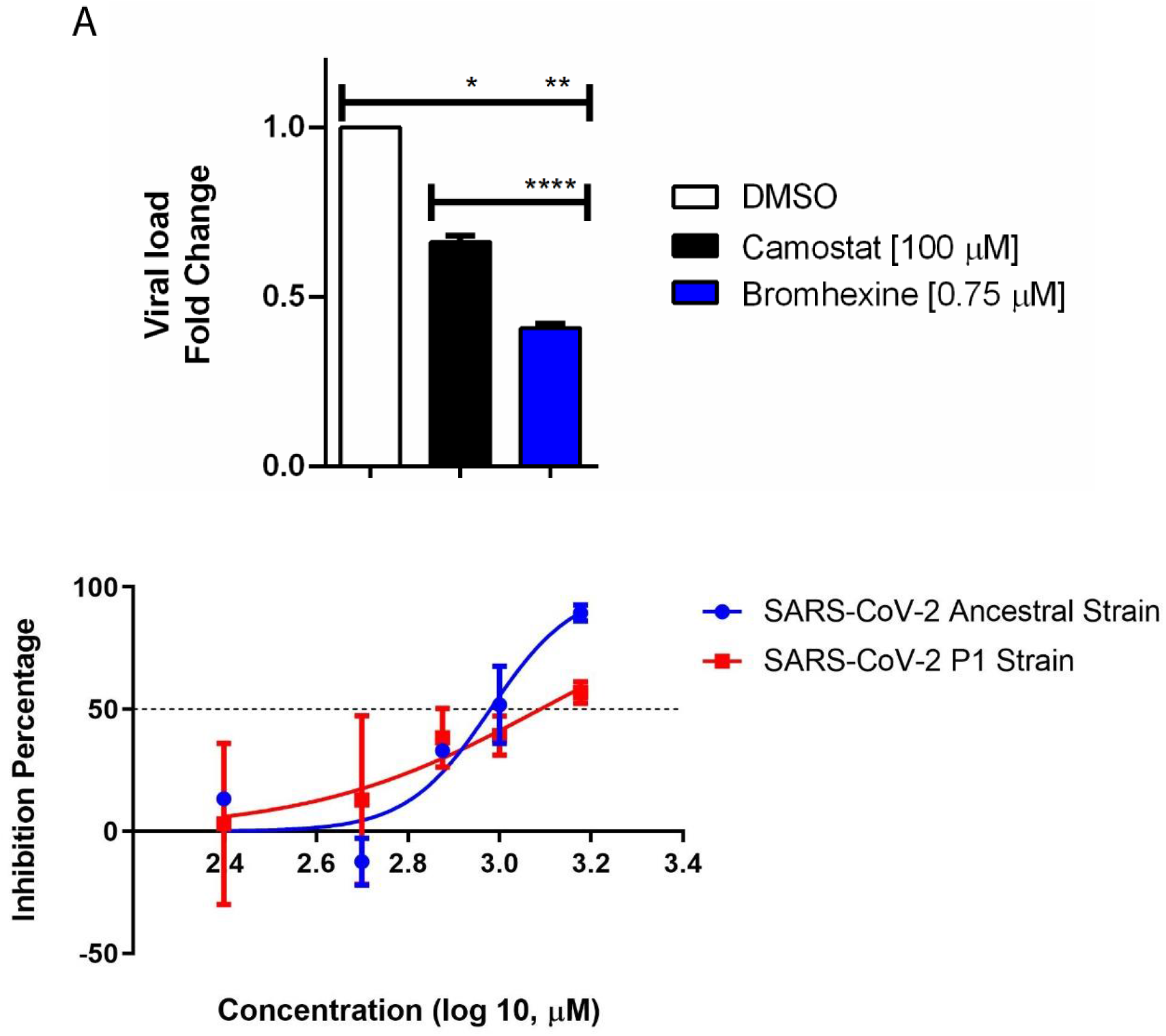
BHH inhibitory effect for SARS-CoV-2 in vitro in Caco-2 cells. A. Absolute quantitation of viral load of SARS CoV-2 in Caco-2 cells treated for 48 hours with BHH at 0.75 μM, Camostat, and mock (DMSO 0.004%); B. Dose–response effect of increasing BHR concentrations in the SARS-CoV-2 RNA progeny production in parental linage (blue line) and P.1 linage (red line) (\**P* < 0.05; \*\**P* < 0.01; \*\*\*\**P* < 0.001).

**Fig 5.**
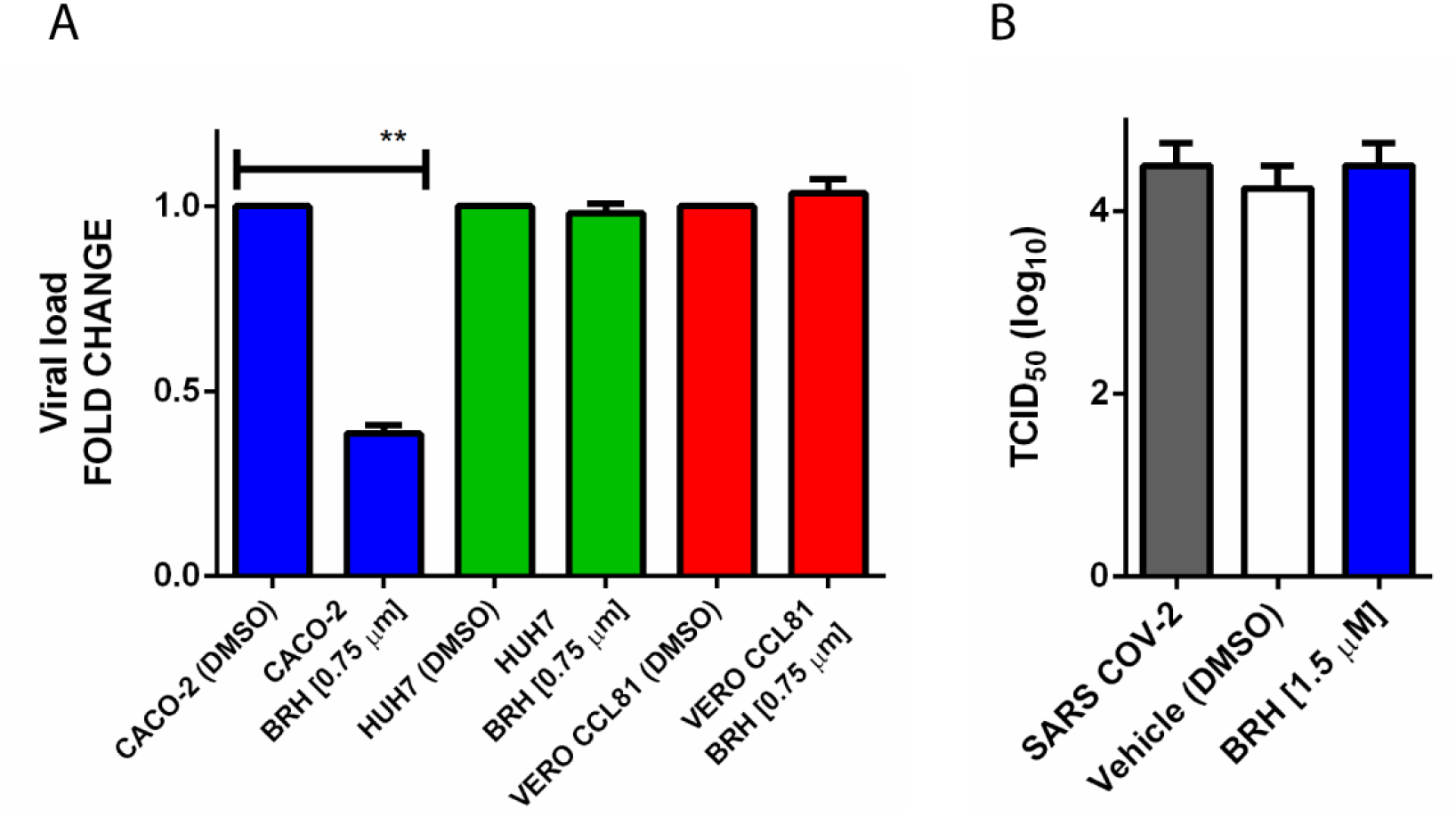
BHH *in vitro* effects over SARS CoV-2 infection. A. BHH replication inhibition to SARS CoV-2 infection of Caco-2, HUH7, and Vero linages. B. Virucidal properties of BHH treatment represented by SARS CoV-2 titers with TCID_50_ assay.

## 4. Discussion

TMPRSS2 was identified as having a critical role in SARS-CoV-2 cell membrane fusion at the cell surface (Shen et al., 2017; Yoshikawa et al., 2019) and is correlated to SARS-CoV-2 cell entry, particularly in the epithelia of lungs and gastrointestinal tract (Mengzhen et al., 2020). With that in mind, inhibition of TMPRSS2 became a potential target against SARS CoV-2 infection. Bromhexine is a mucolytic compound approved by FDA and derived from the alkaloid vasicine, which has been identified as a compound to inhibit TMPRSS2 (Fu et al., 2020; Shen et al., 2017). To study the potential of bromhexine as an antiviral, we tested its activity in SARS-CoV-2 infection in vitro using different cell lineages. We verified that bromhexine significantly reduces the production of SARS-CoV-2 progeny in Caco-2 cells in a dose-dependent manner. The outcome observed with Caco-2, known to express TMPRSS2 (mRNA and protein) (Bertram et al., 2010), was not reproduced with HUH7 and Vero CCL-81 cells. This difference could be explained by a relatively low expression level of ACE2 in hepatocytes (e.g. HUH7) and by differences in the non-human primate TMPRSS2 on Vero CCL-81 cells (Mengzhen et al., 2020). Furthermore, bromhexine showed a higher inhibition effect than camostat, a widely known TMPRSS2 inhibitor.

Additionally, BHH inhibited the replication of VSV-eGFP-SARS-CoV-2-SΔ21 in Caco-2 cells by 40%, which reflects a specific effect on virus-cell binding and entry. However, BHH reduced an average of 90% of the SARS-CoV-2 supernatant progeny in Caco-2 cells. These contrasts may reflect the targeting of TMPRSS2 or other related protease molecules located elsewhere other than the plasma membrane, such as in the Golgi, that may act during the assembly or release of the virus (Clososki et al., 2020). Another conundrum in the present study was the differences in BHH inhibition between SARS CoV-2 WT and gamma (P1) variants. Although BHH effects were significant for both variants, BHH reduced the replication of the gamma variant with lower intensity when compared to WT. The gamma variant has several spike protein mutations that could enhance its avidity for ACE2, with less dependency on TMPRSS2 for cell attachment/entry (Sabino et al., 2021).

BHH had low cytotoxicity for Caco-2 cells (50% toxicity at 76.5 μM), thus being more tolerable than human primary conjunctival fornix epithelial cells (50% toxicity at 45 mM) (Wang et al., 2020). Moreover, TMPRSS2-deficient mice are healthy (Kim et al., 2006), suggesting that the inhibition of TMPRSS2 may not harm the host. Thus, it is conceivable that TMPRSS2 inhibitors, such as bromhexine, could be used to treat SARS-CoV-2 infections.

Results from a clinical trial showed that treatment with 8 mg BHH three times a day for two weeks improved COVID-19 patient outcomes, avoiding ICU admission, assisted ventilation, and mortality (Ansarin et al., 2020). Mikhaylov et al., 2022, showed that 8 mg BHH prophylaxis was associated with a reduced rate of symptomatic COVID-19 patients. Another study revealed that BHH tablets (32 mg) had beneficial effects on COVID-19 patients, with improved chest computerized tomography (CT) and a lesser need for oxygen therapy (Li et al., 2020). The oral use of BHH has been shown to promote mild side effects. Since no substantial pharmacological interactions have been reported, bromhexine can likely be administered with other medicines to obtain a synergistic effect as adjuvant therapy for COVID-19. Recently, in contrast with our data, a study to identify inhibitors of TMPRSS2, done with a biochemical assay using active TMPRSS2 protease and a fluorogenic peptide substrate, showed no inhibition of TMPRSS2 by BHH (Shrimp et al., 2020). Another conflicting observation was described by Hörnich et al., 2021 showing a lack of inhibitory activity of BHH by TMPRSS2 fusion assay. But, in the same study, experiments with SARS-CoV-2 clinical isolate on Calu-3 cells in vitro infection demonstrated significant inhibition of replication by ambroxol and bromhexine; each agrees with our results. Nevertheless, some bromhexine COVID-19 clinical trials showed that early treatment or prophylaxis use of BHH was associated with an 80% improvement in symptomatic COVID-19 patients (Ansarin et al., 2020; Li et al., 2020; Mikhaylov et al., 2022).

Taken together, our findings show that the interaction of BHH with TMPRSS2 active site blocks the protease ability to cleave and activate S, and BHH inhibits SARS-CoV-2 infection and replication in vitro. Our results support the inhibitory activity of BHH during SARS-CoV-2 infection, decreasing about 40% of viral entry and 90% of viral progeny in the supernatant. BHH inhibitory effect may vary depending on the degree of TMPRSS-dependent fusion for different SARS CoV-2 variants. Additional evidence from clinical trials revealed a better prognosis for BHH-treated COVID-19, enhancing that BHH treatment has the potential as an adjuvant drug against coronavirus disease 2019.

## CRediT authorship contribution statement

Eurico Arruda and Priscyla D. Marcato designed the study. Daniel M. M. Jorge performed in silico analyzes. Ronaldo B Martins, Iasmin Ferreira, Leticia Almeida, Juliano P. Souza, Marjorie Pontelli, Italo A. Castro, Thais M. Lima, and Rosa M. M. Viana designed and executed the experimental SARS-CoV-2 infection. All authors contributed to writting the manuscript. Eurico Arruda, Priscyla D. Marcato, and RBM wrote the final version of the manuscript. All authors have approved the final manuscript.

## Acknowledgements

We are thankful to the Brazilian National Council for Scientific and Technological Development (CNPq) (grant number 403201/2020-9), and the São Paulo Research Foundation (FAPESP) (grant number 2019/26119-0). We are grateful to Dr. Edison L. Durigon (Institute of Biomedical Science, University of São Paulo, Brazil) for sharing the SARS CoV-2 clinical strains WT and P1, and to Dr. Sean P.J. Whelan (Department of Molecular Microbiology, Washington University School of Medicine, St. Louis, MO, USA) for sharing the VSV-eGFP-SARS-CoV-2.

